# Temporal Dynamics of EEG Decoding for Continuously Changing Visual Stimuli

**DOI:** 10.64898/2026.01.23.700794

**Authors:** Ilker Duymaz, Micha Engeser, Daniel Kaiser

## Abstract

Multivariate analyses of M/EEG data are typically performed on neural responses time-locked to discrete stimulus onsets. Such designs usually reveal high decoding performance during the initial transient response (0-500 ms), which subsequently drops to a lower, sustained level. Here, we examined time-resolved EEG decoding of natural scene processing when scenes gradually enter the visual field without a clear onset. We created video sequences in which one scene category (e.g., a beach) smoothly transitioned into another category (e.g., a forest) by blending images from two categories into a single composite panorama and moving a square aperture across it. We then compared EEG decoding for the first scenes within the transitions, which appeared with a sudden onset, to the second scenes, which emerged gradually as the videos progressed. For the first scenes, we observed robust category decoding from 60 ms after onset with a clear peak structure. For the second scene, category decoding was markedly weaker and showed no discernable peak structure. Realigning the appearance of category-diagnostic content for the second scene using deep neural networks did not enhance decoding or recover a peak structure. Further, classifiers trained on the first scene generalized to the second, but with a broad, temporally diffuse pattern, indicating that the second scene did not engage the same hierarchical temporal cascade as the first. Together, these results demonstrate that sudden versus gradual onsets produce distinct temporal decoding dynamics. Insights from onset-based decoding studies, therefore, do not straightforwardly extend to continuous and free-flowing natural stimulation.

---

During the last decade, multivariate decoding of magneto-/electroencephalographic (M/EEG) signals has become an essential tool for probing the temporal dynamics of visual representations in the human brain (Grootswagers et al., 2017; King & Dehaene, 2014). M/EEG decoding studies, by and large, employ trial-based designs that treat stimuli as discrete, independent events, separated by blank interstimulus intervals. Stimuli are thus presented with abrupt onsets, where a blank screen is suddenly replaced by the stimulus. When time-locked to these sharp onsets, M/EEG signals typically yield robust decoding shortly after the stimulus onset, with a consistent discernable peak structure within the first 500 ms, followed by a sustained but gradually diminishing trend toward chance-level accuracy even when the stimulus remains visible and consciously perceived (e.g., Carlson et al., 2013; Cichy et al., 2014; Grootswagers et al., 2017; Kaiser, Azzalini, et al., 2016; Kaiser, Oosterhof, et al., 2016).

Many decoding studies draw their primary conclusions from the relative timing of decoding within the first 500 ms of processing, interpreting it as a window into perceptual encoding and category-specific representation (Contini et al., 2017; Wardle & Baker, 2020). For instance, Cichy et al. (2014) contrasted superordinate and subordinate categories, interpreting differences in the timing of onset-locked early peaks to chart hierarchical processing of objects. Likewise, Kaiser et al. (2018) varied object position and observed a robust, onset-locked advantage for typical locations at ∼140 ms, demonstrating that object processing is sensitive to the statistical structure of the environment even at early stages of visual processing. These and many other studies rely on abrupt stimulus onsets and report strictly onset-locked processing latencies. It remains unclear to what extent the nature of these conclusions depends on this specific design choice.

The visual system is highly sensitive to onsets, exhibiting greater detection accuracy (Cole et al., 2004) and enhanced event-related potential (ERP) components (Van Pelt et al., 2025) compared to other visual events such as visual offsets or spontaneous changes in the luminance of already visible stimuli. This advantage has been attributed to the high salience of onsets for attentional capture (Yantis & Jonides, 1984, 1990) and to neural mechanisms tuned to transient, flicker-like cues (Todd & Gelder, 1979; Tolhurst, 1975). From an evolutionary standpoint, this sensitivity may serve the quick detection of predators or environmental hazards that threaten survival. However, in everyday life, our visual experience rarely involves onsets as abrupt as those employed in many visual perception experiments. Instead, we typically encounter gradual transitions, such as when scanning a scene, moving through space, or as objects move through our visual field. This discrepancy raises a central question: to what extent are the temporal structures reported in conventional, onset-locked paradigms a consequence of the abrupt, experiment-imposed onsets that elicit them?

To investigate this question, we developed a paradigm in which one natural scene gradually transitioned into another without a discrete visual onset. To create these transitions, we used generative AI to seamlessly blend images from different scene categories into composite panoramas. We then generated video stimuli by panning a square aperture smoothly across these panoramas, such that the visible content gradually shifted from one scene to the next (Figure 1a). By recording EEG as participants viewed these videos, we aimed to examine how the brain represents visual scenes that emerge gradually, in contrast to scenes that appear abruptly (Figure 1c). This approach allowed us to directly compare neural representations elicited by sudden onsets versus more naturalistic scene transitions.

**Figure 1.**
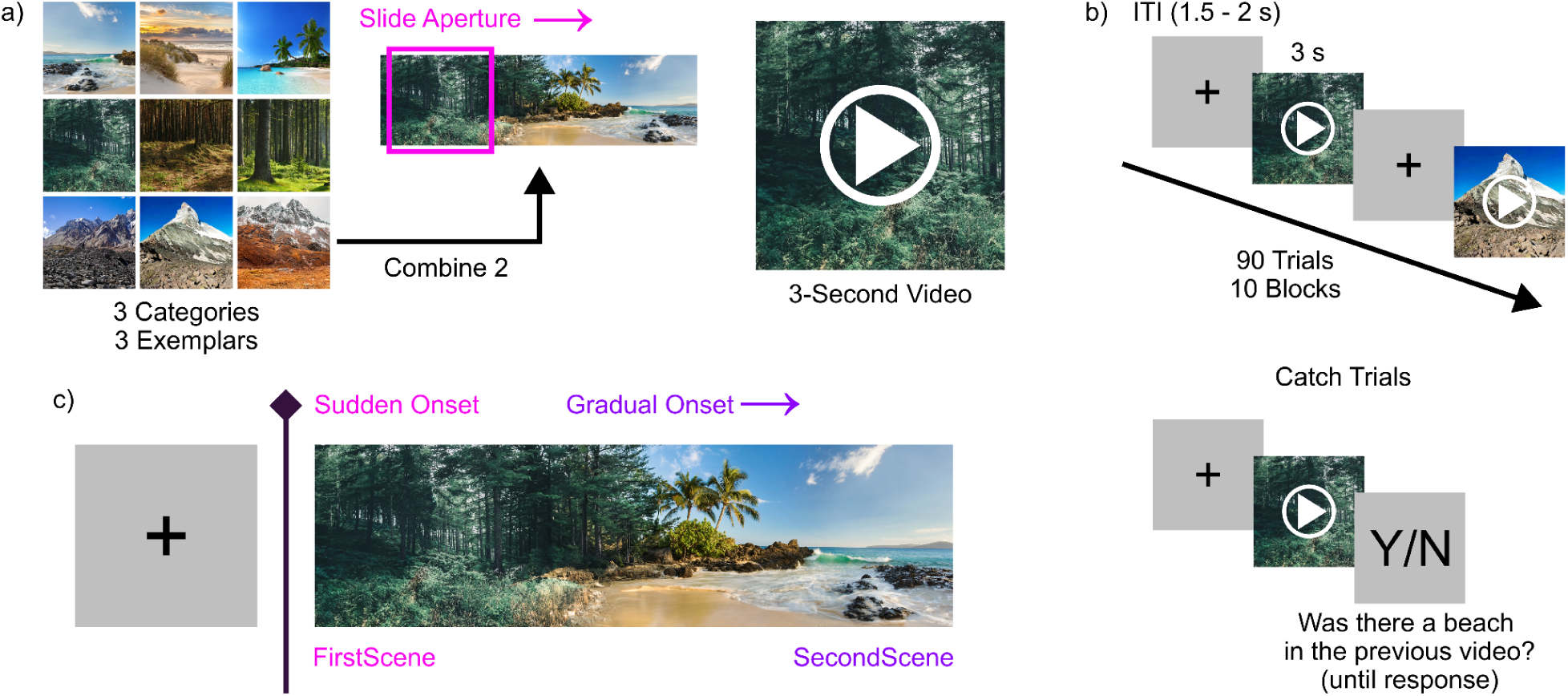
Experimental Design. a) Scene transition videos were created by combining two scenes into a wide panorama and moving a square aperture across it. b) Trial sequence. Each trial began with a 1.5-2 s randomly jittered inter-trial interval, followed by a 3 s transition video, after which the task advanced automatically to the next trial. On a random 20% of trials, a catch prompt appeared after the video, asking whether a specified scene category had been present in the preceding video. c) Since each transition video was preceded by a blank screen, the first scene appeared with an abrupt onset, whereas the second scene gradually entered view as the video progressed.

We found that scenes with sudden onsets exhibit distinct temporal dynamics compared to those with gradual onsets. Sudden onsets produced sharp, transient peaks in decoding accuracy, whereas gradual onsets showed a slower rise to a sustained, and markedly lower, accuracy level. Representations during the transient peak generalized to gradual-onset scene representations in a broad, temporally non-specific pattern, suggesting little to no correspondence between the hierarchical processing of scenes with sudden and gradual onsets. These results point towards pronounced differences that need to be taken into account as we move from trial-based paradigms to more free-flowing stimulation.

## Method

### Participants

35 healthy adults (19 female, 16 male; mean age: 26.9 years, SD = 5.09 years) with normal or corrected-to-normal vision participated in the experiment. Participants provided written informed consent prior to the experiment session and received monetary compensation. The study protocol was approved by the general ethical committee of Justus Liebig University Giessen. All experimental protocols were in accordance with the Declaration of Helsinki.

### Stimuli

The stimuli comprised videos depicting natural scene transitions between three categories: beach, forest, and mountain. For each category, we selected three distinct exemplar images, yielding a total of nine base images. We created composite panoramic scenes by combining all possible pairs of exemplars on a 3500 × 1000 pixel canvas. In each composite, we placed different scene exemplars on the left and right edges of the canvas. We filled the central (initially empty) region using Adobe Photoshop’s (version 25) generative fill tools to blend the two edge images into a naturalistic transition. When needed, we provided prompts to guide the generation, removing artifacts and ensuring physically plausible transitions. We then converted each panoramic image into a 3-second video stimulus (60 frames-per-second), in which a square aperture smoothly panned from the left to the right edge, simulating a naturalistic scene transition (Figure 1a).

We generated 90 unique videos: 81 based on all pairwise combinations of the nine exemplars (9 × 9), and nine based on mirrored versions of identical exemplar pairs (i.e., same-same pairs mirrored after panorama generation). Each condition appeared ten times, resulting in 900 total trials per participant.

### Procedure

We recorded 64-channel EEG while participants viewed the scene transition videos on a 32-inch Gigabyte AORUS FI32Q monitor (resolution: 2560 × 1440 pixels; refresh rate: 120 Hz). Participants sat 60 cm from the screen using a chinrest to maintain consistent viewing distance and minimize head movement. We instructed participants to maintain their gaze on a fixation cross in the center of each video.

Trials began with a fixation cross presented on a gray background for a jittered duration between 1 and 1.5 seconds (in 100 ms increments), followed by the 3-second video subtending 10° × 10° of visual angle on the same gray background (Figure 1b). On 20% of trials, a response prompt appeared immediately after the video, requiring a yes/no response regarding the presence of a specified scene category (e.g., “Was there a mountain in the previous video?”) with equal likelihood for each answer. The prompt remained on screen until response, and no feedback was provided. Participants were performing very well on this task (mean accuracy = 88.5%, SD = 11.7%). On the remaining 80% of trials, the next trial began immediately after the video. Trials were organized into 10 experimental blocks, with participants taking self-paced breaks between blocks.

### EEG Acquisition and Preprocessing

We recorded EEG using a 64-channel ActiCAP system with an ActiCHamp amplifier (Brain Products GmbH, Gilching, Germany) at a 1000 Hz sampling rate. Electrodes were positioned according to the international 10-10 system. Preprocessing and subsequent analyses were performed using MNE-Python (Gramfort et al., 2013). Data were epoched from -1 to 3 seconds relative to stimulus onset and band-stop filtered at 48-52 Hz to remove line noise. We then re-referenced the data to the average across all electrodes, downsampled it to 200 Hz, and applied baseline correction using the -0.5 to 0 seconds pre-stimulus interval. Noisy channels were identified and removed via visual inspection. Blink-related artifacts were removed using Independent Component Analysis (ICA).

### Time-Resolved Decoding

To examine the temporal dynamics of visual representations during sudden versus gradual scene onsets, we trained classifiers at each time point to decode labels corresponding to the first and second scene categories (forest, beach, and mountain) in each transition. Separate classifiers were trained to decode the first- and second-scene category labels from the same time-series data, allowing us to track the evolving representational content in EEG signals as visual stimuli changed over time.

We excluded trials in which the first and second scenes were the same exemplar. In such cases, decoding would yield identical results for both scene labels, making it impossible to distinguish between the representations of the two scenes.

Classification was performed using Linear Discriminant Analysis (LDA) with a time-resolved searchlight approach, allowing us to track changes in decoding accuracy across time. We decoded between three scene categories and applied 10-fold cross-validation with folds balanced at both the category level and the individual video level. Within each fold, the number of trials per category was matched so that the classifier was trained and evaluated on an equivalent amount of data from each category. In addition, trials were partitioned such that repetitions of the same underlying set of scene transitions were distributed across folds (one repetition of each unique scene transition per fold). Classification performance was quantified as accuracy (chance = 33.3%).

To identify time points at which classification performance was significantly above chance, we used a cluster-based permutation test with threshold-free cluster enhancement (TFCE; Smith & Nichols, 2009) as implemented in MNE-Python, generating the null distribution via 10,000 sign-flip permutations (Maris & Oostenveld, 2007).

### DNN-Extracted Scene Onsets

The second scenes in our transitions lacked a discrete onset, as they were gradually blended in from the first scene. As a result, the point at which the second scene became visually recognizable, and thus capable of eliciting a distinct onset response, likely varied across stimuli. This variability in the perceptual emergence of the second scene could smear the temporal decoding profile for the second scenes, given that these scenes would asynchronously pass through the visual hierarchy.

To account for this, we used a deep neural network (DNN) trained on scene recognition (VGG16-Places365; Zhou et al., 2018) to estimate the perceptual emergence of the second scene for each transition individually. For each frame of the transition, we extracted embeddings from the output layer of the DNN, which were 365-dimensional vectors containing probabilities for each scene category included in the Places365 training dataset. Using these probability vectors, we calculated the cosine similarity between each frame and the final frame of the scene transition, which corresponded to the fully revealed second scene. We then fit a sigmoid function to the sequence of cosine similarities across frames to model the emergence of the second scene.

From each fitted sigmoid, we identified three threshold points corresponding to 25%, 50%, and 75% similarity to the second scene as candidate onsets. We used these thresholds to re-align the EEG data, shifting each trial’s time series to align signals based on these DNN-extracted onsets. We then recalculated second-scene decoding accuracy using the aligned data.

This approach allowed us to test whether aligning trials based on DNN-estimated perceptual onsets would reveal a more pronounced decoding peak and improve classification performance compared to analyses using unaligned data.

### Temporal Generalization

We conducted a temporal generalization analysis (King & Dehaene, 2014) to investigate whether neural representations of scenes with sudden onsets (first-scene representations) generalize to those of scenes with gradual onsets (second-scene representations). To this end, we trained classifiers at each time point of the EEG data using the first-scene labels and tested them across all time points using second-scene labels. If second-scene representations share the same neural code as first-scene representations, we would expect a cluster of decoding accuracies significantly above chance when classifiers are trained on the early and tested on the later time points within each trial.

In addition to this cross-decoding analysis, we also examined temporal generalization within each scene type. For these complementary analyses, classifiers were trained and tested across all time points using only first-scene labels, and then separately using only second-scene labels. This enabled us to characterize the temporal stability of representations for each scene type and to assess whether any cross-decoding effects could be explained by the temporal generalization structure observed within each scene. Within-scene temporal generalization matrices were computed using the same preprocessing pipeline, classifier parameters, and cross-validation procedure as in the cross-decoding analysis.

Temporal generalization decoding was performed using a Linear Discriminant Analysis (LDA) classifier across time. As in the time-resolved decoding analyses, a leave-one-out cross-validation approach was used, where each trial was excluded once as a test set, and the classifier was trained on the remaining trials. EEG data were resampled to 100 Hz prior to analysis for computational efficiency. The resulting accuracy matrices (time x time) were smoothed separately for each participant using Gaussian smoothing with a square kernel (SD: 2, kernel size: 9x9 time points).

To assess statistical significance, we again applied cluster-based permutation testing with Threshold-Free Cluster Enhancement (TFCE; 10,000 permutations) to identify time points with decoding accuracy significantly above chance.

## Results

### Time-Resolved Decoding

To examine the temporal dynamics of visual representations under sudden versus gradual scene onsets, we trained classifiers to decode the category labels of both the first and second scenes in each transition.

First-scene category decoding was significantly above chance between 60 ms and 2.06 s, peaking at 41.8% accuracy around 115 ms (Figure 2a). In contrast, second-scene decoding did not show a distinct peak. Instead, decoding became significant after 1.3 s, gradually ramping up without a prominent peak, and reaching a maximum accuracy of 33.6%. This gradual buildup suggests a more temporally diffuse emergence of category information for scenes with gradual onsets.

**Figure 2.**
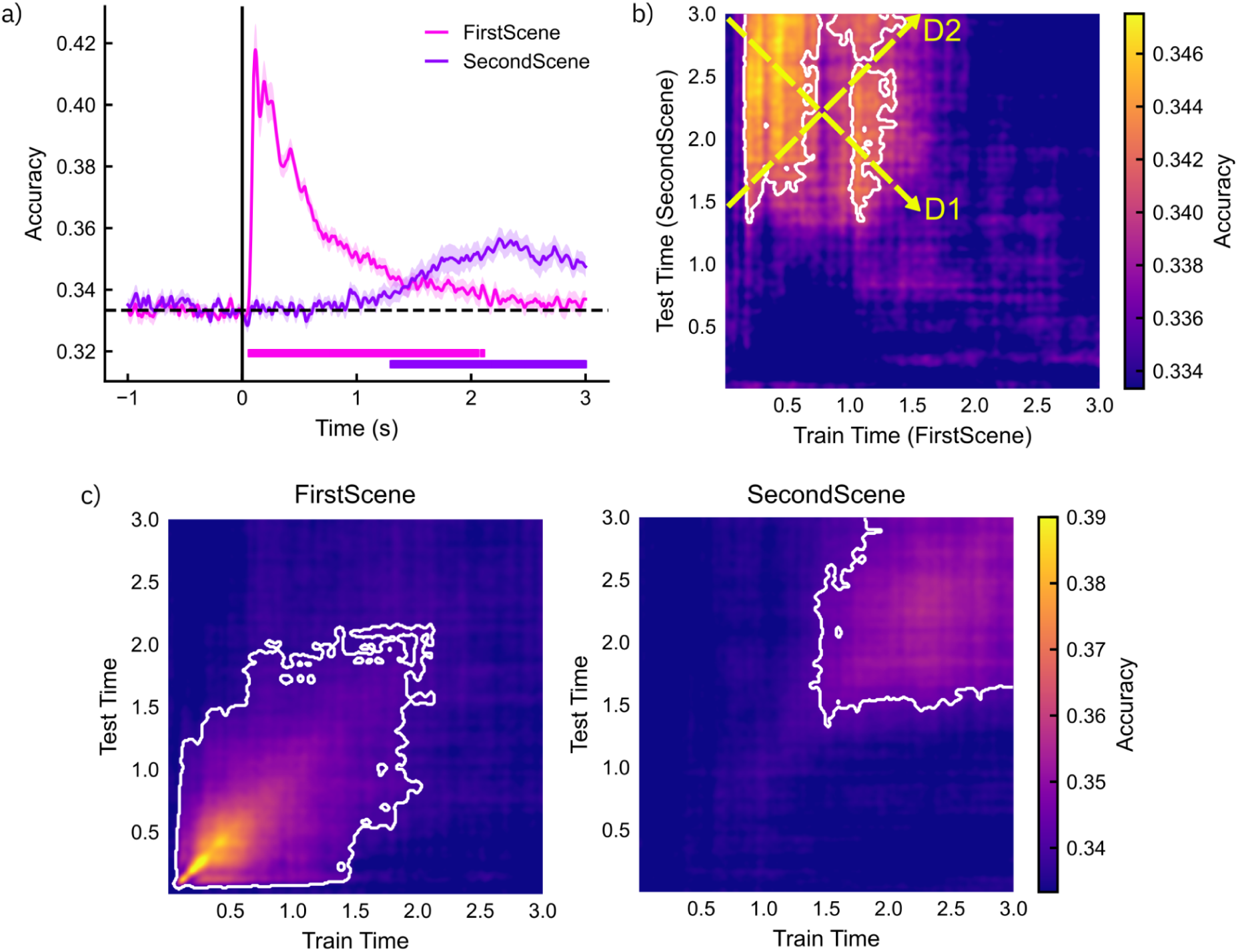
Decoding Results. a) Time-resolved category decoding accuracy for the first scene (sudden onset) and second scene (gradual onset). Category information for the first scene emerged rapidly and yielded multiple discernable peaks, whereas category information for the second scene increased gradually without a distinct peak structure, indicating markedly different temporal dynamics of visual representation under sudden versus gradual onsets. Horizontal bars: p < 0.05. b) Temporal cross-generalization across first- and second-scene category labels. Classifiers were trained using the first-scene labels, and tested on second-scene labels in a separate set of trials, across all timepoints. Temporal generalization revealed a broad, temporally diffuse cross-decoding pattern rather than a forward (D2) or reversed off-diagonal (D1) structure. This indicates that first- and second-scene representations do not share a common hierarchical processing sequence, and that second-scene representations instead stabilize rapidly with minimal temporal evolution. D1: Expected diagonal generalization pattern if the classifier relied on frame-to-frame similarities. D2: Expected diagonal generalization pattern if decoding were driven by correspondence between hierarchical processing stages. Outlined areas: Significant clusters (p < 0.05). c) Temporal generalization analyses for first (left) and second scenes (right). Classifiers were trained on either first- or second-scene labels and tested across all time points in an independent dataset. Decoding for the first scenes was most pronounced along the diagonal of the matrix, consistent with hierarchical processing, whereas decoding for the second scenes was not.

### Temporal Generalization

To investigate whether first- and second-scene representations share common encoding dynamics, we performed a temporal generalization analysis in which classifiers were trained on first-scene category labels at each time point and then tested on second-scene labels across all time points in a separate set of trials. Classifier performance was quantified using decoding accuracy, with a chance level of 1/3 (≈33.3%). Statistical significance was assessed using a Threshold-Free Cluster Enhancement (TFCE; 10,000 permutations).

Overall, there was robust temporal cross-generalization from 150-1350 ms (FirstScene) to 1500-3000 ms (SecondScene), with a brief interruption around the 720-1000 ms interval (p < .05; Figure 2b). Surprisingly, we observed a temporally unspecific, broad generalization pattern, rather than more pronounced decoding along an off-diagonal in the generalization matrix. Given that the first and second scenes in the transition videos were mirrored, and thus time-reversed, versions of one another, a classifier driven primarily by frame-to-frame similarity would therefore be expected to yield a generalization pattern along a reversed off-diagonal (e.g., the first frame of one scene resembling the last frame of the other; Figure 2b, D1). Conversely, if both scenes engage the same hierarchical processing cascade once they appear, their representations would become more similar when corresponding processing stages align in time, yielding the opposite off-diagonal pattern (Figure 2b, D2).

The broad, temporally diffuse generalization we observed instead suggests two possibilities: either the classifier tracked stable representations of scene category throughout the transition, or it relied predominantly on low-level features shared across exemplars (e.g., spatial frequency, color, luminance). In either case, the hierarchical processing order does not appear to correspond between first- and second-scene representations in either a forward or a backward direction. Rather, second-scene representations appear to rapidly reach a relatively stable code, with little evidence of temporal evolution thereafter. This is further supported by the within-scene temporal generalization results, which exhibit a clear diagonal pattern only for first scenes but not for second scenes (Figure 2c).

### DNN-Extracted Scene Onsets

Because the second scenes were blended in gradually rather than appearing with a discrete onset, variability in their emergence likely introduced temporal jitter across transitions, which could smear the onset-locked neural response and obscure a clear peak in second-scene decoding accuracy. Therefore, we realigned signals to DNN-estimated perceptual onsets to test whether this would improve decoding performance and/or recover a peak structure for second-scene decoding. We recalculated second-scene decoding accuracy after shifting trials to each of the three candidate thresholds (25%, 50%, 75% similarity to the second scene). We conducted separate repeated-measures ANOVAs on the average decoding accuracy within the first 500 ms after the DNN-extracted onset, with threshold (unaligned, 25%, 50%, 75%) as a within-subjects factor. We selected the 500 ms time window because we anticipated the strongest initial responses during this interval. This also ensured that offset responses occurring at the end of the videos were excluded from all realigned signals.

The analysis did not reveal a significant main effect of threshold (F(3, 102) = 1.82, p = .148, η²p = .051). None of the aligned conditions produced a qualitatively different temporal decoding pattern, and none yielded a distinct peak structure in second-scene decoding (Figure 3). Instead, the decoding time courses remained comparably diffuse across thresholds. Thus, DNN-based realignment could not recover any resemblance to the sharp onset-related peaks observed for first-scene decoding.

**Figure 3.**
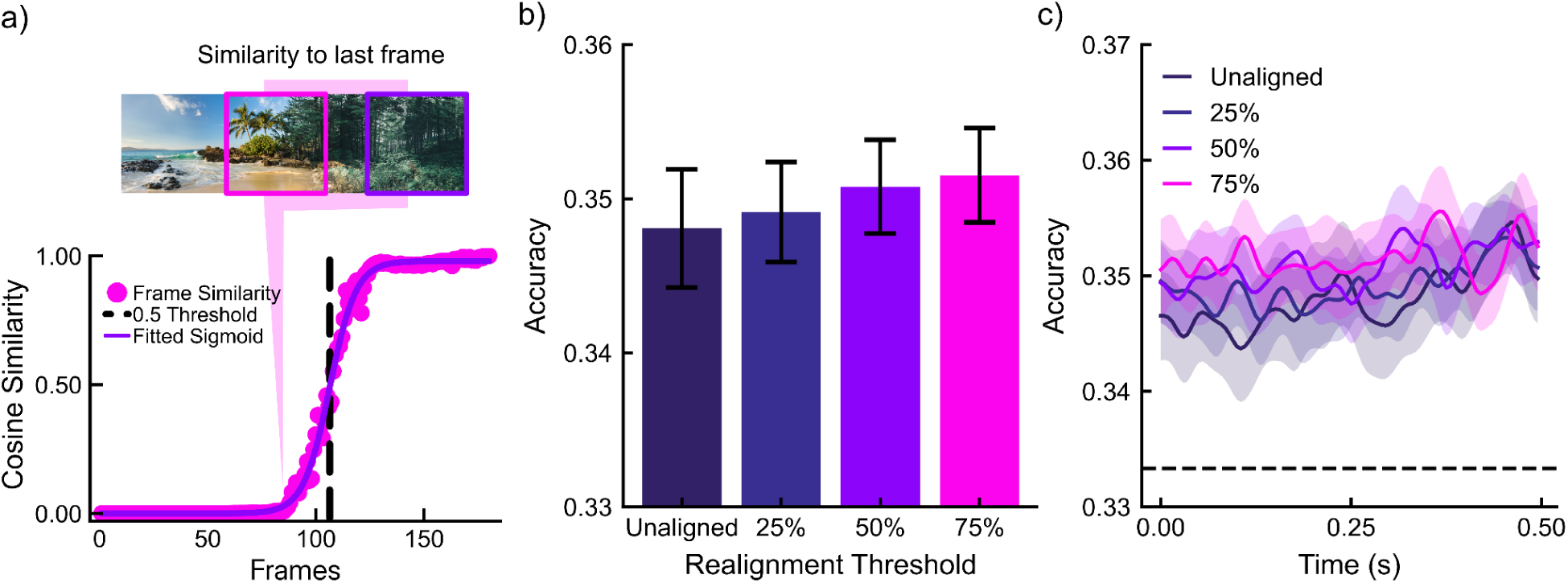
DNN-Extracted Scene Onsets. (a) Example condition showing frame-wise similarity to the second scene (i.e., the last frame of the video; dots) with a fitted sigmoid (line). Candidate onsets are defined as the first frame at which the fit crosses each threshold (25%, 50%, 75%), and trials are realigned to these onset times. (b) Mean second-scene decoding accuracy evaluated for the 500 ms window after the DNN-extracted onset for each alignment condition (25%, 50%, 75%) and for the unaligned baseline (stimulus-onset-locked signals cropped at 1.5-2s). None of the realignment thresholds significantly improved the mean decoding accuracy within the 500 ms time window following the DNN-extracted onsets. Error bars: SEM. (c) Time-resolved decoding accuracy aligned to each threshold (and unaligned baseline). None of the aligned conditions produces a distinct second-scene peak; trajectories remain relatively flat across thresholds, consistent with the absence of a sharp onset-locked response.

## Discussion

We investigated how time-locking to visual onsets in typical trial-based paradigms shapes the dynamics of M/EEG decoding. To this end, we compared EEG decoding for scenes that appeared with sudden, discrete onsets to scenes that emerged gradually during smooth panoramic transitions. We observed three key results. First, sudden onsets reproduced the well-established pattern of a pronounced early decoding peak at approximately 60 ms (peak accuracy = 41.8%), followed by a decay that approached, but remained above, chance (Figure 2a). Second, gradual onsets lacked a discrete peak structure, instead showing a slower ramp to a lower, sustained decoding level (peak accuracy = 33.6%). Third, time-generalization analyses revealed that representations at the onset-locked peak generalized to gradual-onset scenes, indicating a shared representational code despite distinct temporal profiles (Figure 2b). Together, these findings suggest that the widely reported early decoding peak in trial-based designs is strongly shaped by abrupt visual transients, whereas more naturalistic, gradual transitions yield slower-building but sustained category information, implicating design-dependent constraints on how M/EEG decoding indexes visual scene representations.

Unlike classical ERP approaches, which emphasize transient, stimulus-locked peaks, multivariate decoding methods provide a richer characterization of how information unfolds continuously over time. By tracing representational dynamics rather than discrete components, decoding approaches offer a principled way to investigate the seamless evolution of category information as perceptual inputs change. Our results, however, show that the temporal structure of the stimulus strongly shapes these dynamics. In particular, the abrupt onsets widely used in traditional trial-based paradigms yield decoding profiles that diverge substantially from those elicited by gradual onsets, which are arguably more prevalent in, and more representative of, natural vision. This raises the possibility that a narrow focus on early, onset-locked decoding peaks may undermine one of the core advantages of multivariate approaches over ERPs and limit the generalizability of resulting interpretations to real-world visual processing.

Reassuringly, our findings show that representations evoked by sudden onsets generalized robustly to those elicited by gradual onsets. This demonstrates that although abrupt visual onsets modulate the temporal profile and apparent strength of decoding, they do not fundamentally change the underlying representational format. In this sense, the core neural code appears stable across different modes of stimulus presentation. As a result, for studies primarily aimed at establishing whether specific stimulus information is represented in the brain, irrespective of the precise temporal dynamics, the choice between sudden and gradual onsets is unlikely to alter the essential interpretive conclusion.

However, our results also raise the question of how well the dynamics observed in trial-based paradigms capture the neural processes that unfold in the real world, where discrete visual onsets are largely absent. Many M/EEG decoding studies with sudden stimulus onsets have reported latencies for the emergence of representations at different levels (e.g., Cichy et al., 2014, 2016; Grootswagers et al., 2019). While these benchmarks have been valuable for mapping processing under abrupt-transient stimulation, their exact latencies may not translate to naturalistic viewing. In everyday vision, incoming information is temporally correlated, and predictive mechanisms can pre-activate and sustain representations before diagnostic sensory evidence arrives (Battistoni et al., 2017; Peelen et al., 2024). From a predictive-processing perspective, higher-level categorical expectations may at times precede and shape lower-level sensory representations (Battistoni et al., 2017; Peelen et al., 2024), in contrast to the strictly bottom-up timeline implied by onset-locked designs. Consequently, onset-locked latency estimates should be treated as design-contingent markers rather than general laws of visual hierarchy, and our understanding of processing order should be supplemented with paradigms that incorporate continuous, naturalistic input.

At the same time, one might argue that abrupt changes in visual input are not entirely unnatural, because each saccade brings a new region of the scene into foveal vision, effectively creating a sequence of quasi-onset events. Consistent with this, combined eye-tracking and EEG studies have identified robust saccade-locked responses during free viewing, including fixation-related components and category-sensitive effects that closely resemble those reported in classic, fixation-enforced paradigms (e.g., Dimigen & Ehinger, 2021; Gert et al., 2022). Moreover, recent evidence suggests that fixation-related potentials can exhibit faster temporal dynamics than stimulus-onset-locked activity (Auerbach-Asch et al., 2023), and that stimulus-related responses are more tightly aligned with saccade onsets than with subsequent fixation onsets (Amme et al., 2024); consistent with top-down, action-oriented emergence of visual representations that is already underway before the new fixation is established. Together, these findings point to a sophisticated coordination of goal-directed behavior, predictive processing, and peripheral vision in shaping visual representations, and underscore the need for future decoding work that explicitly models saccade-locked dynamics during free viewing to assess how far onset-based paradigms generalize to natural vision. Our results, while most directly informative for paradigms that restrict eye movements, reinforce this need by revealing markedly different temporal dynamics for representations of more naturalistic stimuli.

What causes the marked differences in temporal profiles for sudden versus gradual onsets? Our data do not arbitrate among mechanisms, but we outline three not mutually exclusive accounts. First, abrupt onsets may recruit neurons tuned to transient, flicker-like input (Todd & Gelder, 1979; Tolhurst, 1975), briefly boosting signal-to-noise ratio and producing the sharp decoding peak, with responses then decaying; the absence of which would also align with the lower, sustained accuracy for gradual onsets. Second, visual onsets are highly salient and can trigger bottom-up attentional capture (Yantis & Jonides, 1984, 1990), thereby enhancing visual representations immediately after a sudden appearance. Third, a sudden onset may reset the phase of ongoing oscillations (Hanslmayr et al., 2006; Makeig et al., 2002), synchronizing population activity across trials during the transient window and improving temporal alignment. With the present data, we cannot determine the relative contributions of these mechanisms and therefore do not prioritize one account over the others.

It is worth noting three limitations of our study. First, the perceptual onset of second scenes likely varied across transitions and across repetitions as participants became familiar with the sequences, which can blur time-resolved decoding estimates and flatten transient peaks. We attempted to mitigate this by realigning trials to DNN-estimated perceptual onsets, but residual across-repetition variability may still have contributed to the absence of a distinct peak structure. Second, our stimulus set was relatively small. With only three exemplars per category and correlations among within-category exemplars, classifiers may have overfit to low-level visual features rather than semantic content. As a result, our findings are not well-suited for comparing the temporal dynamics of high-level category representations across onset regimes, although they still elucidate how sudden onsets shape the temporal profile of low-level visual decoding. Third, in our design, the visual system was already adapted to ongoing stimulation by the time the second scenes appeared. This may have reduced overall response magnitudes for the second scene and thereby limited decoding performance. However, our primary interest was in the temporal structure and cross-generalization of decoding rather than absolute signal strength, so adaptation is unlikely to undermine our main conclusions.

Taken together, our results show that sudden visual onsets amplify and sharpen decodable scene information for a brief interval, whereas gradual onsets yield lower-amplitude, and more sustained decoding profiles, even though the underlying representational format generalizes across onset types. These findings imply that onset-locked latency estimates should be treated as design-contingent markers rather than general laws of visual hierarchy, and that inferences about processing order are best supplemented with paradigms that incorporate continuous, naturalistic input. More broadly, our findings highlight that a simple design choice (sudden versus gradual onsets) systematically shapes time-resolved decoding and motivate paradigms that align M/EEG measurements with the temporally continuous structure of everyday vision.

## Acknowledgements

We thank Rico Stecher and the students in the Neural Computation I course for help with experiment setup and data collection. We further thank Malaika Alphonsus, Tanja John, Pietra Pacheco Alves, and Nasibeh Babaei for their assistance in data collection. This work was supported by the DFG (KA4683/5-1; project number 518483074) and under Germany’s Excellence Strategy (EXC 3066/1 “The Adaptive Mind”, project number 533717223). No conflicts of interest, financial or otherwise, are declared by the authors.

## Data Availability

All data and code are publicly available: https://doi.org/10.5281/zenodo.18326081

## Author Contributions

I.D., M.E., and D.K. conceived and designed research; I.D. and M.E. performed experiments; I.D. and D.K. analyzed data; I.D., M.E., and D.K. interpreted results of experiments; I.D. prepared figures; I.D. and D.K. drafted the manuscript; I.D., M.E., and D.K. edited and revised manuscript; I.D., M.E., and D.K. approved final version of manuscript.

